# Thermoadaptation of EndoG proteins in the *Xenopus* frog genus

**DOI:** 10.64898/2026.04.14.718403

**Authors:** Alexander A. Tokmakov

## Abstract

*Xenopus* is a genus of entirely aquatic frogs found in sub-Saharan Africa. Currently, the complete genomes of two species within the *Xenopus* genus, *Xenopus laevis* and *Xenopus tropicalis*, have been fully sequenced, annotated, and made publicly available. The two species inhabit markedly different environments: *X. tropicalis* lives in the hot, equatorial regions of Africa, whereas *X. laevis* resides in the cooler climates of southern Africa. In the present study, mutational profiling, comparative homology modeling, and computational bioinformatics were used to identify the features of adaptive evolution in *Xenopus* endonuclease G (EndoG) proteins. The multiple characteristics of EndoG isozymes were discovered to vary considerably between the two *Xenopus* species dwelling in different locations. Most notably, EndoG proteins from the psychrophilic *X. laevis* exhibit the increased contents of charged and polar residues, elevated pI, higher intramolecular interaction energies, B factors, molecular void volumes, and solvent accessibilities, but the decreased contents of nonpolar and aromatic amino acids, lower hydrophobicity, buried surface area, and molecular packing density compared to those from the thermophilic *X. tropicalis*. The observed differences strongly suggest that temperature plays a dominant role in EndoG diversification. Evaluation of intramolecular interaction energies appears to be a particularly sensitive and discriminative framework for assessing protein divergence at the structural level. Overall, this study highlights the diversification of homologous proteins in ectothermic vertebrate eukaryotes and provides mechanistic insight into protein adaptation to contrasting environments.

## 1. Introduction

The genus *Xenopus* comprises more than twenty species of fully aquatic frogs. The complete genomes of the two species within this genus, *Xenopus laevis* and *Xenopus tropicalis*, have been fully sequenced, annotated, and made publicly available, offering powerful resources for comparative genomic and molecular studies [1,2]. The two species diverged from a common ancestor approximately 34 million years ago [2]. These amphibian species serve as important vertebrate model organisms, yet they have adapted to distinctly different thermal environments over evolutionary time. *X. tropicalis* is native to the hot and humid equatorial regions of sub-Saharan Africa, where it inhabits consistently warm freshwater ecosystems averaging around 26°C. In contrast, *X. laevis* is found in the cooler, more temperate regions of southern Africa, where temperatures average around 20.5°C [3]. In laboratory settings, typical ambient temperatures range 24-28°C for *X. tropicalis* and 17-24°C for *X. laevis* [4-6]. Because amphibians are ectothermic organisms - relying on external environmental sources to regulate their body temperature - their biochemical processes are particularly sensitive to ambient temperature. Given their shared evolutionary background but distinct ecological niches, the two species represent an ideal natural model for studying the molecular mechanisms of thermal adaptation in vertebrate animals.

Protein adaptation to temperature is a well-studied area in molecular biology, biochemistry, and evolutionary biology. Proteins adapt to temperature changes through evolutionary modifications: at elevated temperatures, they evolve increased stability to resist thermal denaturation and maintain their functional structure. Conversely, at lower temperatures, the general slowing of biochemical reactions imposes evolutionary pressure on cold-adapted proteins to acquire increased structural flexibility [7-9]. Major adaptations in thermophilic proteins include enhanced ionic interactions and hydrogen bonding, tight hydrophobic core packing, more rigid structures, and a higher thermal threshold for denaturation. In contrast, psychrophilic proteins are typically characterized by looser hydrophobic cores, fewer stabilizing hydrogen bonds, longer loop regions, and less compact structures [9-11]. Although extensive data have been accumulated concerning the adaptations of thermophilic proteins in bacterial extremophiles [12-15], the adaptations in higher organisms remain underexplored.

EndoG is a nuclear-encoded, nonspecific DNA/RNA nuclease localized to mitochondria in eukaryotic cells. It plays a key role in the regulation of mitochondrial DNA (mtDNA) and has been implicated in apoptosis and other mitochondrial-associated processes. During apoptosis, EndoG is released from mitochondria and translocates into the nucleus, where it contributes to DNA degradation independently of caspase activity [16,17]. Crystal structures of EndoG from several species have been resolved and deposited in the Protein Data Bank [18-20], providing valuable insights into its catalytic and regulatory mechanisms. Structural and biochemical studies have demonstrated that two conserved residues—a histidine and an asparagine—located in the catalytic center play essential roles in magnesium coordination and DNA binding [18,20]. Notably, eukaryotic EndoG contains an N-terminal mitochondrial targeting sequence (MTS) that directs its import into mitochondria [16]. Furthermore, EndoG has been shown to form a catalytically active homodimer, as revealed by both crystallographic and biochemical analyses [18,20]. Multiple EndoG homologs have been identified across different species and cell types, suggesting evolutionary diversification and potential subfunctionalization of this nuclease.

Here, we report that EndoG isozymes differ substantially between the two *Xenopus* frog species *Xenopus laevis* and *Xenopus tropicalis*, which inhabit distinct environmental regions. The contents of charged, polar, hydrophobic, and aromatic amino acids, isoelectric point (pI), hydrophobicity, solvent accessibility, molecular packing density and void volume, B factors, protein stability, and intramolecular interaction energies were all affected by environmental adaptation. The detailed analysis of intramolecular energies revealed distinct contributions from electrostatic, torsional, and hydrophobic components, providing additional insights into the mechanisms of adaptive protein evolution. The observed energetic adjustments in *Xenopus* EndoG proteins appear to counteract thermal destabilization under warmer environmental conditions. Importantly, the results from energy-based calculations closely align with those obtained through alternative structural analyses, such as the estimation of solvent accessibility, packing density, and protein flexibility.

## 2. Methods

### 2.1. Sequence retrieval and alignments

Amino acid sequences of *X. laevis* EndoG (AAH87366.1 and XP_041428654.1) and *X. tropicalis* EndoG (NP_001017202.1 and KAE8584272.1) were retrieved from the NCBI protein database [https://www.ncbi.nlm.nih.gov/]. The annotated amino acid sequence of mouse EndoG (accession number NP_031957.1) was used as a query in a BLASTsearch against the *X. tropicalis and X. laevis* protein databases [https://blast.ncbi.nlm.nih.gov/Blast.cgi]. Sequence alignments were performed using the EMBL-EBI resource CLUSTAL Omega [21; https://www.ebi.ac.uk/jdispatcher/msa/clustalo] and CLUSTALW, hosted on the GenomeNet platform [22; https://www.genome.jp/tools-bin/clustalw]. Phylogenetic analysis was conducted based on the multiple sequence alignment of *Xenopus* EndoG proteins generated with CLUSTALW.

### 2.2. Calculation and prediction of physicochemical parameters

The grand average of hydropathicity (GRAVY) index and isoelectric point (pI) were calculated using the ProtParam tool, while the proportion of buried residues and solvent accessibility were assessed with the ProtScale, both freely available on the Expasy server [23; https://web.expasy.org/protparam/].

### 2.3. Homology modeling

The three-dimensional structures of *Xenopus* EndoG isozymes were built by homology modeling based on the crystal structure of a mammalian homolog. The crystal structure of DNA-free mouse EndoG at 2.80 Å resolution was used as a template to generate the 3D models of *Xenopus* [18]. The coordinate file of the template was retrieved from the Protein Data Bank (PDB: 6LYF). Homology modeling was performed using the SWISS-MODEL protein structure homology modeling server [24,25; http://swissmodel.expasy.org/]. Visualization of the modeled EndoG structures was carried out with DeepView/Swiss-PdbViewer [24; https://spdbv.unil.ch/] and commercial molecular graphics software Waals (Altif Laboratories, Tokyo, Japan). Model quality was validated using Ramachandran plots and QMEAN analysis [26,27]. Superposition of protein structures and calculations of the root mean square deviation (RMSD) were performed using Waals. Computations of intramolecular interaction energies were carried out with the GROMOS96 implementation within the DeepView/Swiss-PdbViewer.

### 2.4. Other methods

Packing parameters of the modeled *Xenopus* EndoG structures, such as molecular volumes and packing densities, were evaluated using the online calculation tool ProteinVolume 1.3 [28; https://gmlab.bio.rpi.edu/PVolume.php]. The parameters of protein flexibility, including B factors and Lindemann coefficients, were evaluated using FlexServ, a web-based tool for analyzing coarse-grained protein dynamics [29; https://mmb.irbbarcelona.org/FlexServ/].

## 3. Results

### 3.1. Mutational profiling in Xenopus EndoG proteins

Several homologous protein sequences were identified by the BLAST search in *X. laevis* and *X. tropicalis* species using the published sequence of mouse EndoG protein as a reference. Each of them had the BLAST score exceeding 200 and covered a full-length protein, which comprised approximately 290 amino acids. The identity of these sequences as EndoG homologs was confirmed through UniProt. The sequences were aligned using ClustalW, and the resulting multiple alignment is presented in Fig. S1A. In total, 38 homologous and non-homologous substitutions and 1 three-amino acids deletion were identified among the sequences by the alignment. A phylogenetic tree of *Xenopus* EndoG proteins was constructed based on the full-length amino acid sequences (Fig. S1B). The sequences cluster into two distinct evolutionary groups, with two closely related EndoG proteins found in each *Xenopus* species, suggesting a gene duplication event that predated the divergence of these species.

All *Xenopus* EndoG proteins share a common structural organization comprising two highly conserved functional domains separated by a long, unstructured interdomain linker and flanked by a short C-terminal patch (Fig. 1A). The essential catalytic residues within the nuclease core (H138 and N169 in NP_001017202.1) are strictly conserved across all proteins. Mapping the identified amino acid substitutions onto the schematic representation of *Xenopus* EndoG revealed a distinct pattern: while the conserved domains exhibit a relatively low frequency of mutations, the interdomain linker displays a substantially higher rate of variation (Fig. 1B). This non-uniform distribution suggests differential evolutionary constraints across the protein, with strong purifying selection acting on structurally and functionally indispensable regions, and relaxed selective pressure permitting greater variability in the flexible linker.

**Fig. 1.**
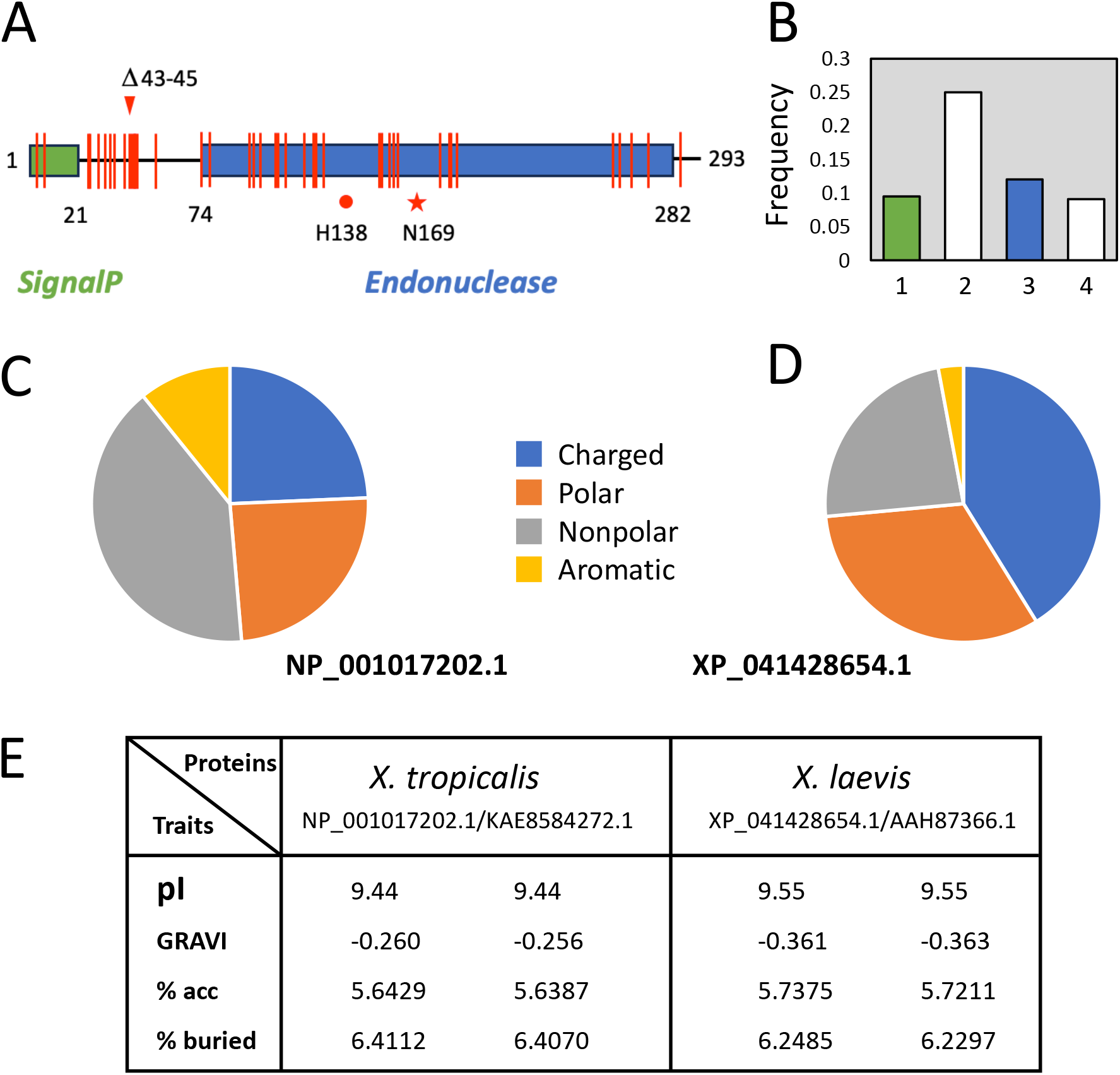
Compositional differences and physicochemical properties of *Xenopus* EndoG proteins. (A) Schematic representation of EndoG protein domains and features annotated according to UniProt, with mapped multiple substitutions in *Xenopus* EndoG proteins. (B) Frequency of substitutions in the signaling domain (1), interdomain linker (2), endonuclease domain (3), and C-terminal tail (4). (C, D) Amino acid composition of substitution sites in *X. tropicalis* and *X. laevis*, respectively. (E) Integral physicochemical properties of *Xenopus* EndoG proteins.

A detailed analysis of substitution sites in EndoG proteins from the two *Xenopus* species revealed clear differences in the physicochemical properties of the affected residues. In *X. laevis*, substitution sites more frequently contained polar and positively charged amino acids, whereas in *X. tropicalis*, nonpolar and aromatic residues were more commonly found (Fig. 1C and D). These compositional differences translated into distinct integral physicochemical properties of the EndoG proteins. Specifically, *X. laevis* EndoG proteins exhibited lower hydrophobicity and a reduced proportion of buried residues, but higher solvent accessibility and isoelectric point (pI), as compared to *X. tropicalis* (Fig. 1E). The potential connection between these molecular properties and the differing environmental conditions of each species’ habitat is discussed further in this paper.

### 3.2. 3D structures of Xenopus EndoG proteins

To access the potential impact of multiple amino acid substitutions within the conserved catalytic core of *Xenopus* EndoG proteins, structural homology modeling was performed for all identified isoforms using the SWISS MODEL server. The crystal structure of mouse EndoG (PDB entry, 6lyf), which shares over 70% sequence identity and aligns with approximately 74% of *Xenopus* EndoG, served as the template for homology modeling. Notably, the largely unstructured N-terminal region (first 62 amino acids) and the short C-terminal tail (last 15 residues) were truncated in the deposited crystal structure of the template protein, and, therefore, could not be modeled using this approach. The quality of the resulting models was validated using QMEAN and GMQE scores, as well as Ramachandran plots (Fig. S2), all of which confirmed proper folding and structural integrity of the models. The generated 3D structure of NP_001017202.1 protein, which covers the entire catalytic core of the protein but lacks the largely disordered N-terminal region and C-terminal tail, is presented in Fig. 3A.

Importantly, despite multiple mutations in the catalytic endonuclease domain (Fig. 1A), its overall structure is fully conserved in all *Xenopus* EndoG proteins. Spatial alignment of the modeled protein molecules from the two *Xenopus* species shows that their structures superimpose remarkably well, with an RMSD of just 0.032 Å across all 218 residues of the molecular core (Fig. 2B). Also, the quaternary assembly of the *Xenopus* EndoG proteins is not affected by multiple substitutions in the two *Xenopus* species. EndoG monomers dimerize with their active sites facing opposite directions, as presented in Fig. 2C. As reported previously, the largely unstructured N-terminal regions of the monomers protrude deep into the catalytic core of the opposite subunits, whereas the C-terminal patches are positioned at the intermonomer interface, supporting the proposed role of these regions in dimer formation [20]. While the modeled 3D structures failed to reveal substantial structural differences between EndoG proteins from the two *Xenopus* species, they enabled comparative analysis of the structure-based protein properties, such as intramolecular interaction energies and packing parameters of monomers and dimers, as presented in the following sections.

**Fig. 2.**
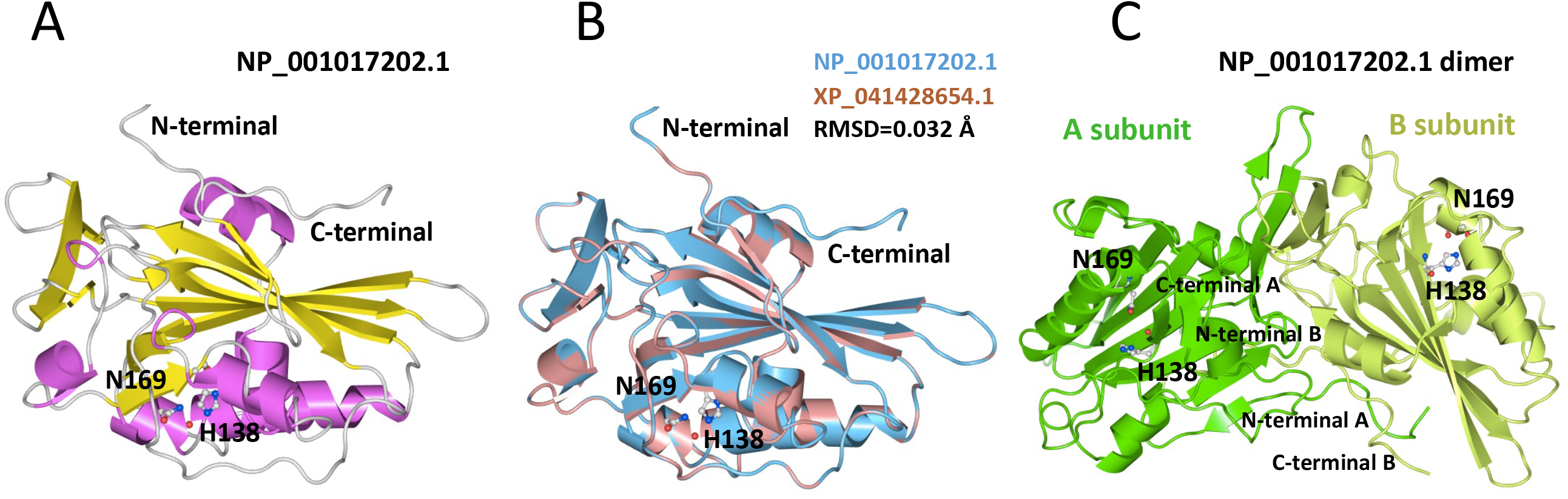
Structural homology modeling of *Xenopus* EndoG proteins. (A) Cartoon representation of the modeled 3D structure of the *X. tropicalis* EndoG protein (NP_001017202.1). (B) Structural superimposition of EndoG proteins from *X. tropicalis* (magenta) and *X. laevis* (red brown). (C) Cartoon representation of the modeled *Xenopus* EndoG dimer. The DNA- and magnesium-binding catalytic center of the enzyme is defined by the conservative catalytic residues H138 and N169.

**Fig. 3.**
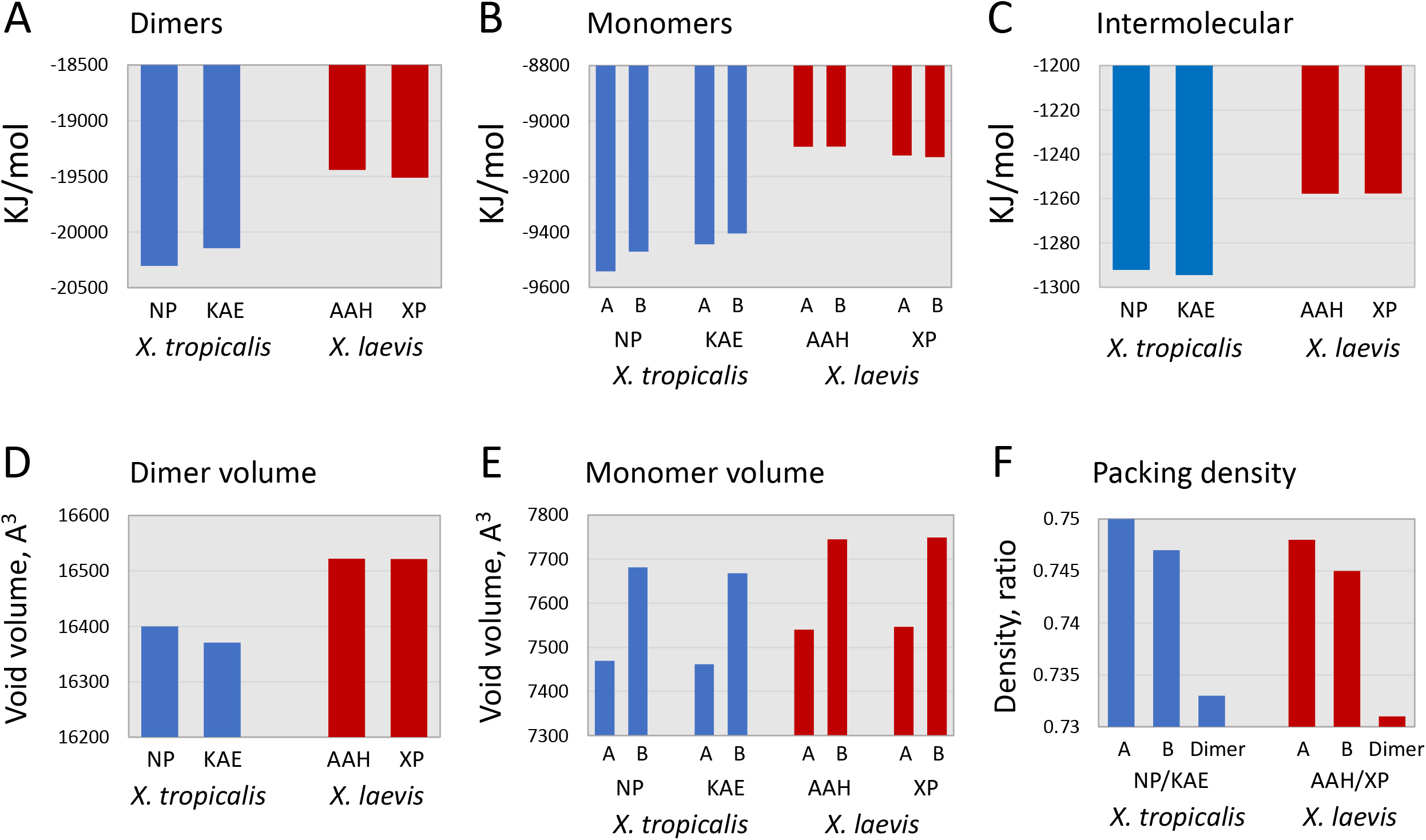
Intramolecular interaction energies and molecular packing of *Xenopus* EndoG proteins. (A, B) Total interaction energies of EndoG dimers and monomers, respectively. (C) Intermonomer interaction energies within EndoG dimers. (D, E) Intramolecular void volumes in EndoG dimers and monomers, respectively. (F) Molecular packing densities of *Xenopus* EndoG monomers and dimers.

### 3.3. Interaction energies in Xenopus EndoG molecules

The interactions within protein molecules play a crucial role in protein folding and stability. A comparative analysis of total interaction energies in *Xenopus* EndoG proteins - calculated from individual amino acid contributions - revealed a distinct difference in the energy state of proteins from the two species. The total interaction energies in EndoG dimers and monomers were found to be lower in *X. tropicalis* than in *X. laevis* (Fig. 3A and B), indicating stronger interactions and potentially higher stability of *X. tropicalis* EndoG proteins. For both dimers and monomers, the difference in total interaction energies amounted to approximately 4%. Importantly, only the energy of electrostatic interactions was substantially lower - by about 7.5% - in *X. tropicalis* EndoG dimers, while other energy components, such as bonds, angles, torsion, improper, and non-bound, showed minimal differences or were slightly higher (Table 1). Combined with the data of mutational profiling (Fig. 1C and D), these findings suggest that the charged residues, that are found in abundance at the substitution sites in *X. laevis* EndoG proteins, contribute to destabilizing the EndoG structure.

**Table 1.** Interac-on energies in *Xenopus* EndoG proteins.

| PROTEIN | bonds | angles | torsion | improper | nonBonded | electrostatic | TOTAL |
| --- | --- | --- | --- | --- | --- | --- | --- |
| *Xenopus tropicalis* |  |  |  |  |  |  |  |
| **NP_001017202.1** |  |  |  |  |  |  |  |
|  | 2179 | 4041 | 2768 | 391 | -14259 | -15425 | -20304 |
| **KAE8584272.1** |  |  |  |  |  |  |  |
|  | 2226 | 4001 | 2779 | 392 | -14175 | -15369 | -21146 |
| *Xenopus laevis* |  |  |  |  |  |  |  |
| **XP_041428654.1** |  |  |  |  |  |  |  |
|  | 2037 | 3932 | 2728 | 360 | -14290 | -14279 | -19512 |
| **AAH87366.1** |  |  |  |  |  |  |  |
|  | 2030 | 3928 | 2771 | 357 | -14282 | -14246 | -19443 |

Note: Interaction energies are given in units of kilojoules per mole (kJ/mol).

Of particular interest is the observation that intermonomeric interaction energy in EndoG dimers differs between the two *Xenopus* species. Specifically, the total energy of intermonomeric contacts is approximately 2.7% lower in *X. tropicalis* than in *X. laevis* (Fig. 3C), suggesting stronger monomer–monomer interactions and potentially greater dimer stability in *X. tropicalis*. Notably, although intermonomeric interactions contribute only a small fraction—approximately 6.5%—of the total interaction energy in the EndoG dimer (Fig. 3A and C), they are still sufficient to render dimer formation energetically favorable.

### 3.4. Packing density and flexibility of Xenopus EndoG molecules

The performed analysis of interaction energies revealed species-specific differences in the capacity of *Xenopus* EndoG molecules to engage in intra- and intermolecular interactions. It was reasonable to hypothesize that these differences would also be reflected in distinct protein packing characteristics and conformational flexibility. Specifically, the lower intramolecular energies and stronger interactions observed in *X. tropicalis* EndoG, suggested a more tightly packed protein structure compared to *X. laevis*. Consistent with this, the void volume in *X. tropicalis* EndoG dimers was smaller than that in *X. laevis* (Fig. 3D). A similar trend was also evident at the monomer level, where both monomers A and B from *X. tropicalis* exhibited lower void volumes than their *X. laevis* counterparts (Fig. 3E). Accordingly, the packing density of both EndoG monomers and dimers was higher in *X. tropicalis* than in *X. laevis* (Fig. 3F). These findings align well with the results obtained by analysis of interaction energies in *Xenopus* EndoG molecules (Fig. 3A-C).

Independent analysis using coarse-grained protein dynamics demonstrated that molecular flexibility is greater in *X. laevis* EndoG compared to *X. tropicalis*, as indicated by the cumulative B-factor—a measure of spatial uncertainty in atomic positions (Fig. 4A). Furthermore, the computed Lindemann coefficients, which quantify the root-mean-square vibrational amplitude of residues, also showed small-scale differences between the two species (Fig. 4B). Particularly, the coefficients for buried residues were consistently higher in *X. laevis* EndoG proteins, pointing to their increased internal mobility. The findings from coarse-grained protein dynamics support the conclusions drawn from the evaluations of interaction energy and packing density. Specifically, the lower interaction energies, increased void volume, and reduced packing density observed in *X. laevis* EndoG are consistent with its enhanced molecular flexibility. Collectively, these structural features point to a less compact and more dynamic conformation of EndoG in *X. laevis* compared to *X. tropicalis*.

**Fig. 4.**
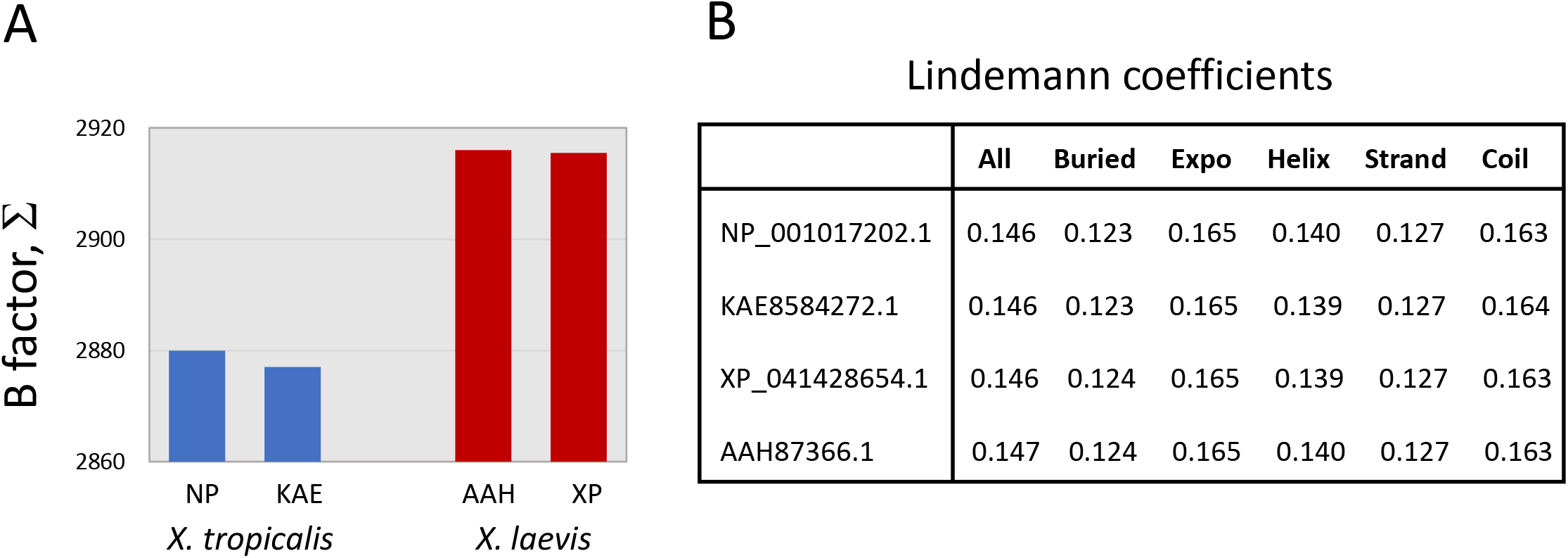
Molecular flexibility of *Xenopus* EndoG proteins. (A) Cumulative B-factors and (B) Lindemann coefficients calculated using coarse-grained protein dynamics.

## 4. Discussion

Early findings on temperature-driven protein evolution have largely been obtained from prokaryotic and invertebrate organisms. Our study extends these principles to vertebrate animals and offers new insights into molecular adaptation in cold-versus warm-adapted species. While previous studies, for example on mollusc cytosolic malate dehydrogenase and abalone enzymes, have compared orthologs from different genera of invertebrate taxa adapted to distinct thermal environments [30,31], our work is among the first to examine protein adaptation in two vertebrate eukaryotic species of the same genus, providing insights into thermal evolution within a vertebrate lineage.

The major findings of this study are consistent with the notion that thermophilic proteins require increased stability to maintain their functional activity at elevated temperatures. Our data confirm that EndoG proteins from the thermophilic species *X. tropicalis* are more stable, compact, and rigid than their counterparts from the cold-adapted *X. laevis* (Fig. 3 and Fig. 4). In addition, *X. tropicalis* EndoG exhibits greater hydrophobicity and buried surface area, along with reduced solvent accessibility, indicating tighter hydrophobic core packing (Fig. 1E). This structural trait represents a well-recognized hallmark of thermophilic protein adaptation. Importantly, the differences extend beyond the monomeric subunits to the dimeric assembly. Not only intramolecular, but also intermolecular interaction energies are lower in *X. tropicalis* EndoG (Fig. 3C), indicating stronger intersubunit cohesion between monomers in the dimeric structure of this thermophilic EndoG. Accordingly, the void volume is lower and packing density is higher in EndoG proteins from *X. tropicalis*, consistent with their tighter spatial packing (Fig. 3D-F). To our knowledge, these findings provide the first clear evidence of environmentally driven adaptation in the quaternary structure of a multimeric protein. Previous studies reported inconsistent patterns of thermal adaptation in the quaternary structure, arguing that oligomerization order plays only a minor role in adaptation to hyperthermophily in bacteria [32].

Previously, it has been demonstrated that amino acid composition shows some correlation with protein thermostability. In general, proteins from thermophilic organisms tend to contain higher proportions of proline and charged amino acids, which contribute to increased structural rigidity and promote favorable electrostatic interactions [10,11]. Charged residues, in particular, often form salt bridges that reinforce protein stability [13, 32]. However, in the present study, an opposite trend emerged: *X. laevis*, a psychrophilic, cold-adapted species, exhibits a higher content of charged residues in its EndoG protein compared to the thermophilic *X. tropicalis* (Fig. 1C and D). Paradoxically, however, electrostatic interaction energy is higher in *X. laevis* EndoG (Table 1). These findings suggest that rather than stabilizing the structure, the excess charged residues in the psychrophilic EndoG promote structural destabilization, thereby enhancing conformational flexibility and catalytic activity at low temperatures. This uncommon adaptive strategy challenges the traditional view of charged residues as stabilizing agents and warrants further investigation.

Notably, the substitution sites in *X. laevis* EndoGs are enriched in positively charged amino acids, resulting in an increased protein pI (Fig. 1E). We hypothesize that this divergence between *X. laevis* and *X. tropicalis* EndoGs may also involve adaptive changes associated with their distinct habitats. However, these substitutions might reflect adaptation to environmental pH rather than thermoadaptation. Indeed, regional variation in water pH across Africa is well documented: in Central Africa, pH ranges from 5.3 to 7.8, whereas in Southern Africa, it varies between 6.79 and 8.36 [33]. It is therefore reasonable to propose that, to maintain high catalytic efficiency, *X. laevis* EndoG has evolved substitutions that shift protein pI toward more alkaline values, consistent with the more alkaline waters of Southern Africa. Previously, adaptation of protein pI patterns to environmental conditions has been reported in bacterial proteomes. In particular, it was observed that the proteomes of intracellular parasites tend to be more alkaline, reflecting their adaptation to the elevated pH of the host environment [34]. In line with this concept, accumulating evidence indicates that protein pH dependence is tuned to subcellular pH. It was found that the average pH of maximal protein stability within a given subcellular compartment correlates with the local pH of that compartment [35-37]. Consistent with these observations, a strong correlation between averaged protein pI and intra-compartmental pH has been reported for the human proteome [38].

Finally, in this study, the intra- and inter-molecular interaction energies, classified into the energies of bonds, angles, torsion, improper, non-bound, and electrostatic in SWISS PROT implementation of the GROMACS algorithm emerged as the most sensitive and discriminative parameters of adaptive evolution in *Xenopus* EndoG proteins (Fig. 3 and Table 1). These energies can be reliably computed from 3D protein structures. Importantly, the results from energy-based calculations closely align with those obtained through alternative structural analyses, such as the estimation of solvent accessibilities, molecular void volumes, packing densities, and B-factors. These integral properties of protein molecules are largely defined by the underlying intramolecular interaction energy landscape. To our knowledge, the evaluation of individual interaction energy components has not been systematically applied in the previous studies of protein evolution. It appears that this approach may provide a general highly sensitive and discriminative framework for assessing protein adaptation at the structural level.

## Data availability statement

The data that support the findings of this study are available within the article and its supplementary materials.

## CRediT authorship contribution statement

Author confirms the sole responsibility for the conception of the study, presented results and manuscript preparation.

## Appendix A.

### Supplementary data

Supplementary data to this article can be found online.

