## Supplemental materials for "Thermoadaptation of EndoG proteins in the *Xenopus* frog genus"

#### A *Multiple alignment*

```
NP_001017202.1      MGFRGRVLLSGVSLAVGAALGAGVAAWRSRDGAGNVGVLDILGFQTVQASTQLSLPPSAG      60
KAE8584272.1        MGLRGRVLLSGVSLAVGAALGAGVTVWRSRDGAGNVGVLDILGFQTVQASTQLSLPPSAG      60
AAH87366.1          MGLRGRALLSGVSLAVGAALGAGVTAWRKRDEAANAGVLRI---PAVQASTQLALPPSAG      57
XP_041428654.1      MGLRGRALLSGVSLAVGAALGAGVTAWRKRDEAANAGVLRI---PAVQASTQLALPPSAG      57
                    **:***.*****************:.**.** *.*.*** *    :*******:******

NP_001017202.1      LTRFGLPGLSQLKSRESYVLSYDPRLRGPAWVLEHLSPERLHGSAERQGCDFQEDVSVHQ      120
KAE8584272.1        LTRFGLPGLSQLKSRESYVLSYDPRLRGPAWVLEHLSPERLHGSAERQGCDFQEDVSVHQ      120
AAH87366.1          LTRFGLPGLSQLKTRESHVLSYDPRLRGPAWVLEHLTPDRLKGSAERKDCEFQEDVSVHH      117
XP_041428654.1      LTRFGLPGLSQLKTRESHVLSYDPRLRGPAWVLEHLTPDRLKGSAERKDCEFQEDVSVHH      117
                    *************:***:******************:*:**:*****:.*:********:

NP_001017202.1      YHRAANSDFKGSGFDRGHLAAAANHKWSQKAMDETFILSNIYPQNPHLNQKAWNNLERYC      180
KAE8584272.1        YHRAANSDFKGSGFDRGHLAAAANHKWSQKAMDETFILSNIYPQNPHLNQKAWNNLERYC      180
AAH87366.1          YHRSTNSDYKGSGFDRGHLAAAANHKWSQKAMEDTFMLTNVYPQNPHLNQKAWNNLEKYC      177
XP_041428654.1      YHRSTNSDYKGSGFDRGHLAAAANHKWSQKAMEDTFMLTNVYPQNPHLNQKAWNNLEKYC      177
                    ***::***:***********************::**:*:*:****************:**

NP_001017202.1      RSLTKKNKNVYVCTGPLFLPRREPDGNMYVKYQVIGSNNVAVPTHFFKVVVLEKFSGEIE      240
KAE8584272.1        RSVTKKNKNVYVCTGPLFLPRREPDGNMYVKYQVIGSNNVAVPTHFFKVVVLEKFSGEIE      240
AAH87366.1          RGLTKSNKNVYVCTGPLFLPRREPDGNMYVKYQVIGSNNVAVPTHFFKVVVLEKFSGEIE      237
XP_041428654.1      RGLTKSNKNVYVCTGPLFLPRREPDGNMYVKYQVIGSNNVAVPTHFFKVVVLEKFSGEIE      237
                    *.:**.******************************************************

NP_001017202.1      LRSYVMPNHPVDEQIPLERFLVPIESIERAAGLLFVPNILKNTNNLKAITAGR      293
KAE8584272.1        LRSYVMPNHPVDEQIPLERFLVPIESIERAAGLLFVPNILKNTNNLKAITAGR      293
AAH87366.1          LRSYVMPNHPVDEQTPLDRFLVPIESIERSAGLLFVPNILKNTSNLKAITAGR      290
XP_041428654.1      LRSYVMPNHPVDEQTPLDRFLVSIESIERSAGLLFVPNILKNTSNLKAITAGR      290
                    ************** **:**** ******:*************.*********
```

#### B *Phylogenetic tree*

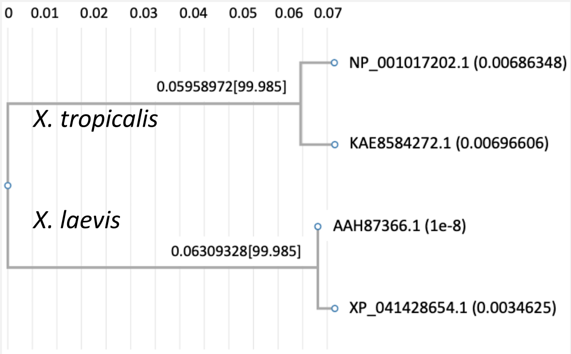


Figure S1

### Validation of generated *Xenopus* EndoG models using QMEAN analysis and Ramachandran plots

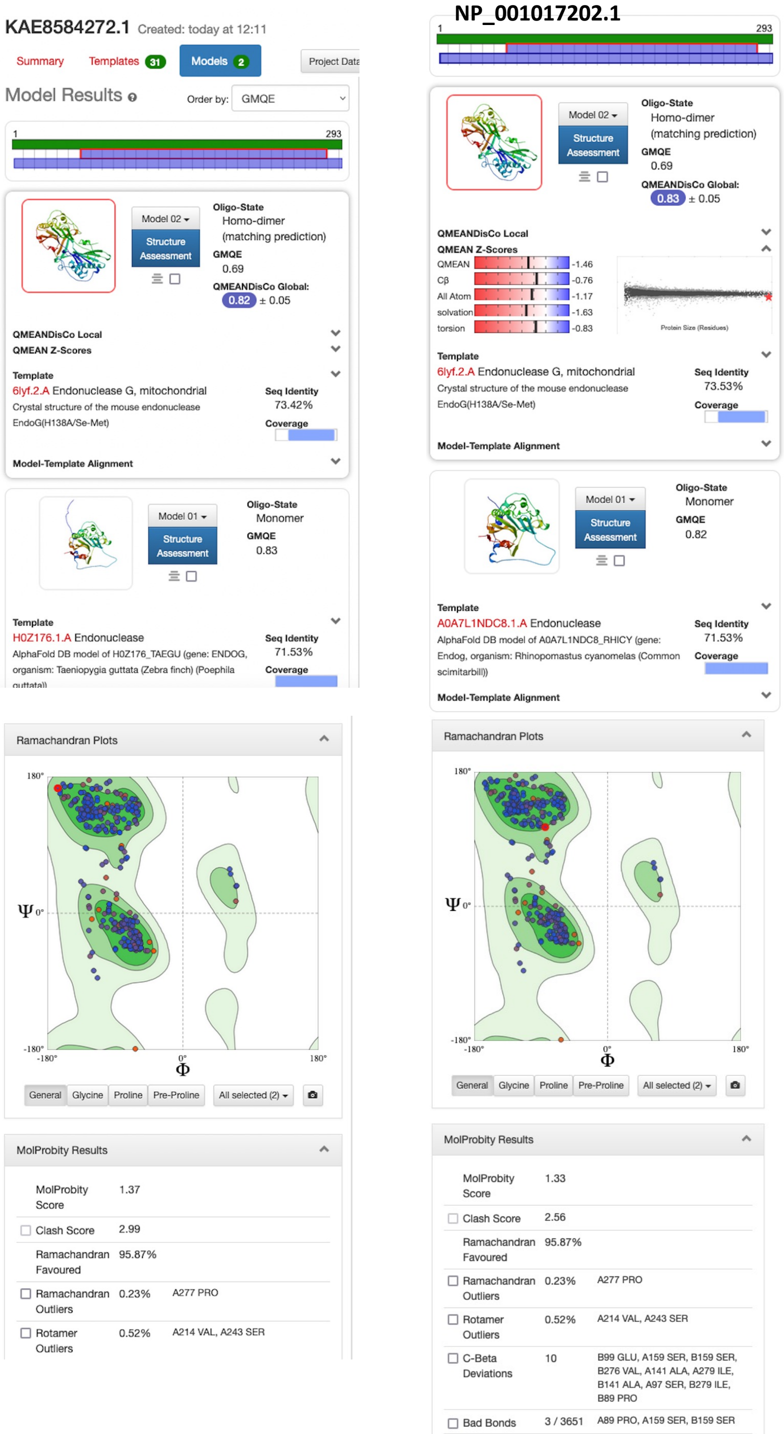


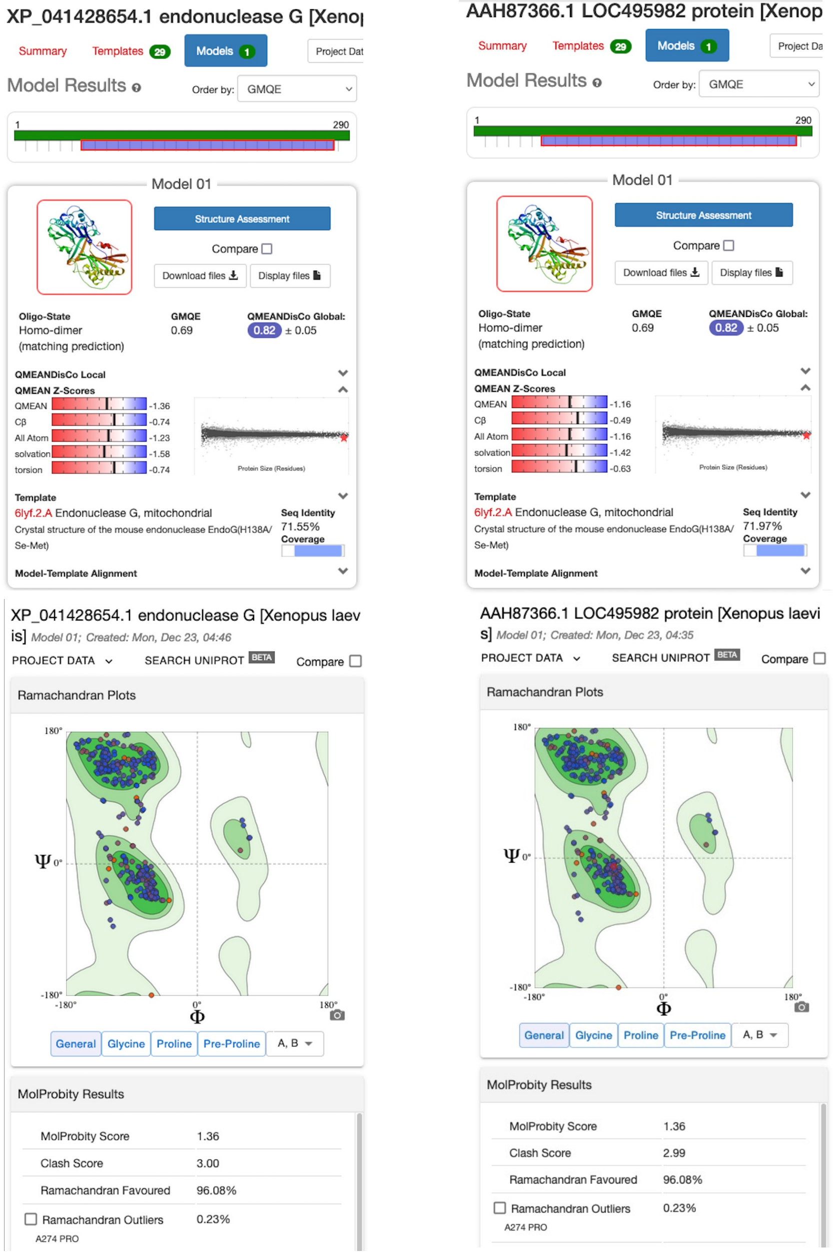


Figure S2
